# A C/ebpα isoform-specific differentiation program in primary myelocytes

**DOI:** 10.1101/2023.05.16.540903

**Authors:** Maria-Paz Garcia-Cuellar, Selin Akan, Robert K. Slany

## Abstract

The transcription factor CCAAT-enhancer binding factor alpha (C/ebpα) is a master controller of myeloid differentiation that is expressed as long (p42) and short (p30) isoform. Mutations within the *CEBPA* gene selectively deleting p42 are frequent in human acute myeloid leukemia. Here we investigated the individual genomics and transcriptomics of p42 and p30. Both proteins bound to identical sites across the genome. For most targets, they induced a highly similar transcriptional response with the exception of a few isoform-specific genes. Amongst those we identified early growth response 1 (*Egr1*) and tribbles 1 (*Trib1*) as key targets selectively induced by p42 that are also underrepresented in *CEBPA*-mutated AML. Egr1 executed a program of myeloid differentiation and growth arrest. Oppositely, Trib1 established a negative feedback loop through activation of Erk1/2 kinase thus placing differentiation under control of signaling. Unexpectedly, differentiation elicited either by removal of an oncogenic input or by G-CSF did not peruse C/ebpα as mediator but rather directly affected the cell cycle core by upregulation of p21/p27 inhibitors. This points to functions downstream of C/ebpα as intersection point where transforming and differentiation stimuli converge and this finding offers a new perspective for therapeutic intervention.

## Introduction

A block of hematopoietic differentiation is a hallmark of myeloid leukemia ^1^. While gain-of-function oncogenes have been intensely studied, less is known about the details of genetic control elements that are perturbed by loss-of-function mutations in AML. An example is the transcription factor C/ebpα that is frequently mutated in myeloid malignancies ^2^. Normal C/ebpα plays a dual role. It cooperates with other transcription factors, including HOX-homeobox proteins, to establish a hematopoietic enhancer landscape necessary for the development of myeloid precursor cells ^3, 4^. Consequently, leukemic transformation through HOX mediated pathways is not possible in the absence of C/ebpα ^5^. On the other side, it is also essential for differentiation. Forced overexpression of C/ebpα causes terminal maturation. This dualism is reflected by the presence of two C/ebpα isoforms. By use of alternative translation initiation codons, either a long p42 isoform associated with differentiation, or a shorter p30 version connected to proliferation can be produced ^6^. Both retain DNA binding functionality but they differ in the extent and activity of an N-terminal transactivation domain. Interestingly, biallelic *CEBPA* mutations in AML are common. However, in all cases at least one allele capable of producing p30 is retained ^7, 8^. In mouse models, specific deletion of p42 causes fully penetrant leukemia ^9^. This has been interpreted as an oncogenic gain of p30 activity and hitherto studies mostly concentrated on the genetic network downstream of p30 ^10–12^. Here, we investigated C/ebpα isotype-specific genomics and transcriptomics in primary hematopoietic precursor cells and its relation to HoxA9 mediated transformation. Our findings suggest a particular role for p42 in establishing a signaling-dependent differentiation program. The oncogenic activity of HoxA9 perturbs this program downstream of C/ebpα itself. Our results emphasize the importance of a loss-of-function for leukemogenesis and give potential perspectives how to bypass this defect for therapeutic purposes.

## Methods

### DNA, cells, inhibitors, antibodies

Retroviral plasmids were constructed in pMSCV (Clontech, Palo Alto, CA) vectors. All insert sequences were either derived from laboratory stocks or amplified from cDNA isolated from murine cells and confirmed by sequencing. Degron constructs were adapted to a murine environment by PCR-based introduction of a F36V mutation into the FKBP moiety ^13^. HPSCs were isolated from C57/BL6 mice with a triple-ko for *Elane*, *Prtn3*, and *Ctsg* ^14^. Transduction was done with CD117 (Kit) selected cells enriched with magnetic beads (Miltenyi, Bergisch-Gladbach, Germany) essentially as recommended by the manufacturer. To generate transformed lines, cells were cultivated in methylcellulose (M3534, StemCellTechnologies, Cologne, Germany) for two rounds under antibiotics selection, then explanted and maintained in RPMI1640 (Thermo-Scientifc, Germany) supplemented with 10% FCS, penicillin-streptomycin, 5ng/ml recombinant murine IL-3, IL-6, GM-CSF, and 50ng/ml recombinant murine SCF (Miltenyi, Bergisch-Gladbach, Germany). dTAG13 was from Tocris (NobleParkNorth, Australia). All other chemicals were provided either by Sigma (Taufkirchen, Germany) or Roth (Karlsruhe, Germany). Antibodies were purchased either from Thermo Scientific, Darmstadt, Germany) or from Cell Signaling Technologies (Leiden, Netherlands).

### ChIP-Seq, cell lysis, nascent-RNA isolation

ChIP was performed as described in ^15^ applying a 10 min crosslink in 1% formaldehyde @ RT followed by lysis in deoxycholate buffer (50mM Tris/HCl pH8.0, 10mM EDTA, 100mM NaCl, 1mM EGTA, 0.1% sodium-deoxycholate, 0.5% N-lauroylsarcosine 1mM PMSF and 1% HALT complete protease inhibitor cocktail (Pierce, Thermo-Fisher, Germany). Precipitation for all samples was performed with protein G coupled paramagnetic beads (Cell Signaling Technologies). Antibodies used for ChIP: anti-HA rabbit monoclonal, Cell Signaling Technologies (#3724) 5µl per 5x106 cells; anti-C/ebpα rabbit monoclonal, Cell Signaling Technologies (#8178) 5µl per 5x106 cells.

Cell lysis for western was done in 20mM HEPES pH 7.5, 10mM KCl, 0.5mM EDTA, 0.1% triton-X100 and 10% glycerol supplemented with 1mM PMSF and 1% HALT complete protease inhibitor (triton lysis) or in hot (95°C) 50mM TrisHCl pH6.8, 0.2% SDS followed by a 2min nucleic acid digestion at RT with 10 units of benzonase after supplementation with 0.5mM MgCl2 (SDS lysis). Nascent-RNA isolation was done exactly as described in ^16^.

### NGS and bioinformatics

ChIP sequencing libraries were prepared using NEBNext® Ultra™ II DNA Library Prep Kit reagents (NEB, Ipswitch, MA) according to the procedure recommended by the manufacturer. Size selection was done after final PCR amplification with Illumina index primers for 14 cycles. Nascent RNA was converted into Illumina compatible libraries with NEBNext® Single Cell/Low Input RNA Library Prep reagents according to the standard protocol. Sequencing was done at the in house core facility yielding 100bp single- or paired-end reads. Data were mapped with BWA mem (0.7.17) ^17^ to the *Mus musculus* mm10 genome. Reads mapping more than once were excluded by filtering for sequences with a mapping quality score > 4. For visualization BAM files were normalized and converted to TDF format with IGV-tools of the IGV browser package ^18^. Peak finding, motif analysis and peak annotation was done with Homer (4.9.1) ^19^. BAM files were converted to bigwig by Deeptools (3.0.0, bamCoverage) ^20^. Metagene plots were created with Deeptools (3.0.0). Matrices were calculated with calculateMatrix and plotted with plotHeatmap from the Deeptools suite. RNA derived reads were aligned with STAR (v020201) ^21^ to the reference genome mm10 and reads derived from repetitive sequences were excluded by samtools (view)1.8 ^22^ . Transcripts were quantified by Homer analyzeRNA routines and further analyzed with standard spreadsheet tools.

### Data availability

Raw NGS reads were submitted to the European Nucleotide Archive under accession number PRJEB862028

### Statistics

Where appropriate two-tailed T-test statistics were applied.

## Results

### Isoform specific expression in primary hematopoietic precursors

The wt *Cebpa* cDNA is preceded by a short upstream reading frame that affects the choice of the start codon within the main coding sequence. To achieve isotype-specific expression and detection we replaced this feature with an optimized Kozak initiation site and supplied an N-terminal HA-tag. This allowed exclusive production of p42 without generation of additional p30 protein (figure 1A). In transient luciferase assays, transactivation capacity of the modified p42 was retained while, as expected, p30 was largely transcriptionally inactive in this setting (supplemental figure 1A). Preliminary experiments showed that we could not achieve stable expression of p42 in hematopoietic precursors due to strong induction of terminal differentiation. Therefore, we additionally modified the constructs with a C-terminal FKBP degron moiety to enable a “stealth” approach (figure 1B). In the presence of the small molecule dTAG, FKBP modified proteins are dimerized with and degraded by the endogenous E3-ubiquitin ligase cereblon. Target cells grown in dTAG can be retrovirally transduced with the respective construct and protein expression is initiated after release from degradation by removing the dimerizer. We also noticed that p42 is selectively cleaved by myeloid granule proteases (predominantly by cathepsinG) after cell lysis (supplemental figure 1 B). As we have described before for HoxA9 and Meis1 ^23, 24^ that are similarly sensitive to granule proteases, this precludes efficient chromatin immunoprecipitation. Therefore all experiments were done in cells harvested from mice with a triple knockout of elastase, proteinase 3, and cathepsin G (*Elane*, *Prtn3*, *Ctsg* triple k.o.) ^14^. As these animals have no hematological abnormalities, there is no indication that this gene deficiency affects normal blood development.

**Figure 1:**
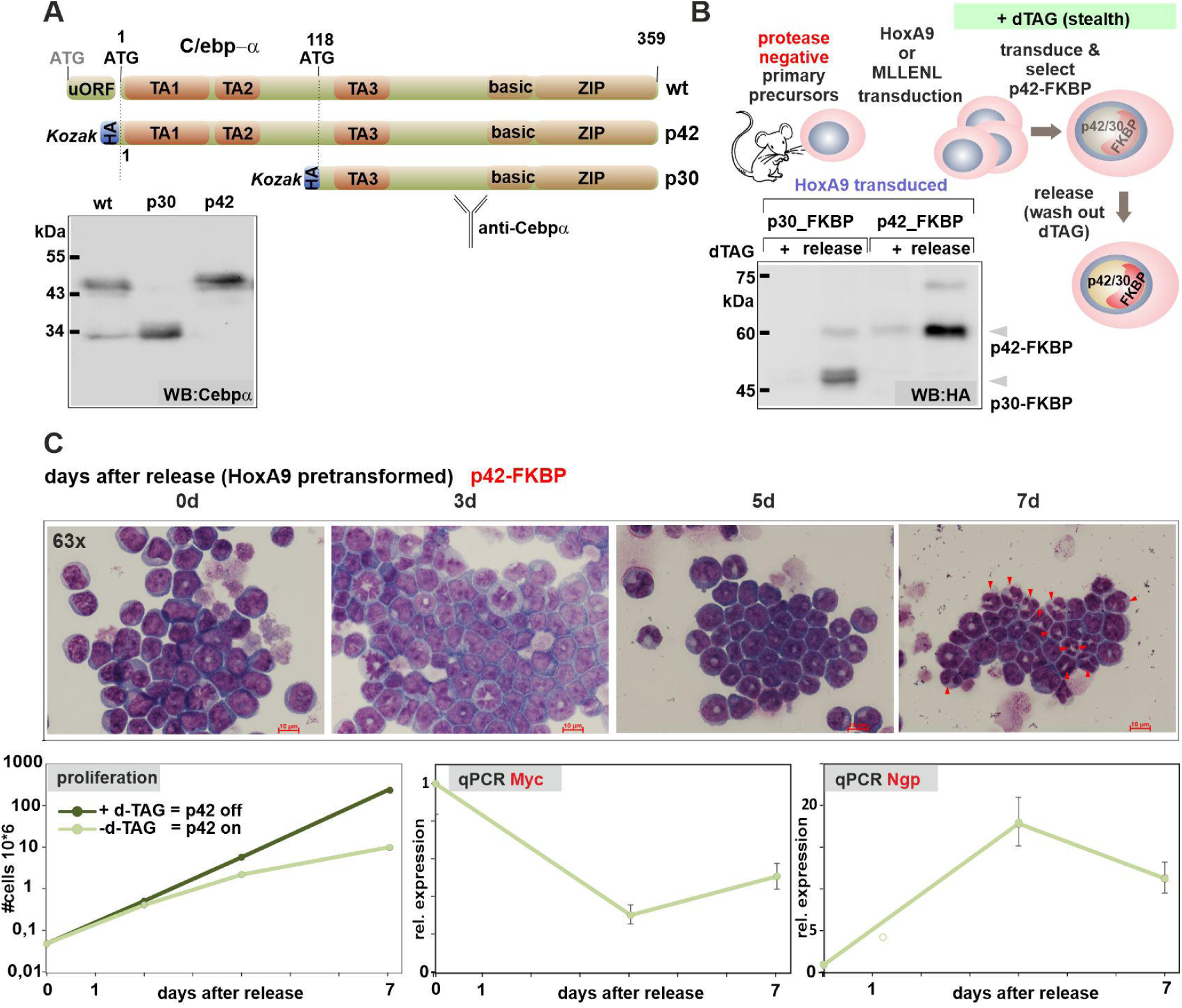
Isoform specific expression of C/ebpα. A: Schematic depiction of wt *Cebpa* gene configuration and changes introduced to achieve isotype specific expression. A short upstream reading frame (uORF) controls use of ATG start codons within the main coding sequence leading to expression of long (p42) and short (p30) C/ebpα isoforms. Replacement of the uORF by an optimized Kozak site allows isoform specific expression as shown in the western blot. B: Experimental set up of inducible C/ebpα expression in HSPCs. Hematopoietic precursors isolated from animals with a knock-out of neutrophilic proteases (*Elane*, *Prtn3*, *Ctsg* triple ko) were immortalized either by HoxA9 or MLLENL and subsequently transduced in the presence of the degrader dTAG with p42-FKBP or p30-FKBP constructs. Expression of the respective proteins was tested in the presence of dTAG and 24h after release from degradation by western blot. C: p42-FKBP is biologically active. HoxA9 x p42-FKBP cells were cultivated for the indicated time in medium without dTAG and morphological aspect, proliferation rates, as well as expression of *Myc* and the differentiation sentinel gene *Ngp* (neutrophilic granule protein) were followed by May-Grünwald-Giemsa staining, counting and qPCR respectively. Micrographs were taken on a Zeiss Axioskop with a Nikon camera at 63x magnification. The size bar corresponds to 10µM.

Induction of p42 expression in primary hematopoietic cells pre-immortalized with HoxA9 induced strong morphological differentiation. This was accompanied by growth arrest and downregulation of *Myc* as well as induction of the differentiation sentinel gene *Ngp* coding for neutrophilic granule protein (figure 1 C). In a similar setting, p30 was mostly inactive (supplemental figure 1C) with only a minor induction of myeloid maturation still observed after expression of p30. To conduct the following experiments independently in two different cell types, we introduced the p42/p30-FKBP constructs also in protease-negative HSPCs pre-immortalized with a MLLENL oncogene (supplemental figure 1D).

### p42 and p30 colocalize on chromatin

Genome wide binding sites specific for p42 and p30 were determined by ChIP with anti-HA antibodies 24h after release of C/ebpα production. Additionally ChIP for endogenous C/ebpα with an antibody recognizing both isoforms was done in immortalized parental cells before transduction with p42-FKBP or p30-FKBP (figure 2A). All binding profiles were highly superimposable for all binding events observed (figure 2B). Across the genome, p42 occupied 33067 sites in HoxA9- and 53098 sites in MLLENL cells that were congruent with p30 and total-C/ebpα peaks (figure 2C). A global analysis revealed a remarkable correlation between p42 occupation density and endogenous C/ebpα as well as with p30 binding. Spearman correlation coefficients of p42/p30 pairs reached 0.91 in HoxA9 and 0.85 in MLLENL cells respectively. Values in this range are usually seen only for direct technical replicates, indicating that p42 and p30 bind to identical sites on chromatin. We could also confirm colocalization of p42/p30 with HoxA9 (supplemental figure 2A). Finally, we verified the known dimerization of p42 and p30 by bidirectional co-immunoprecipitation as biochemical correlate for the colocalization on chromatin (supplemental figure 2B).

**Figure 2:**
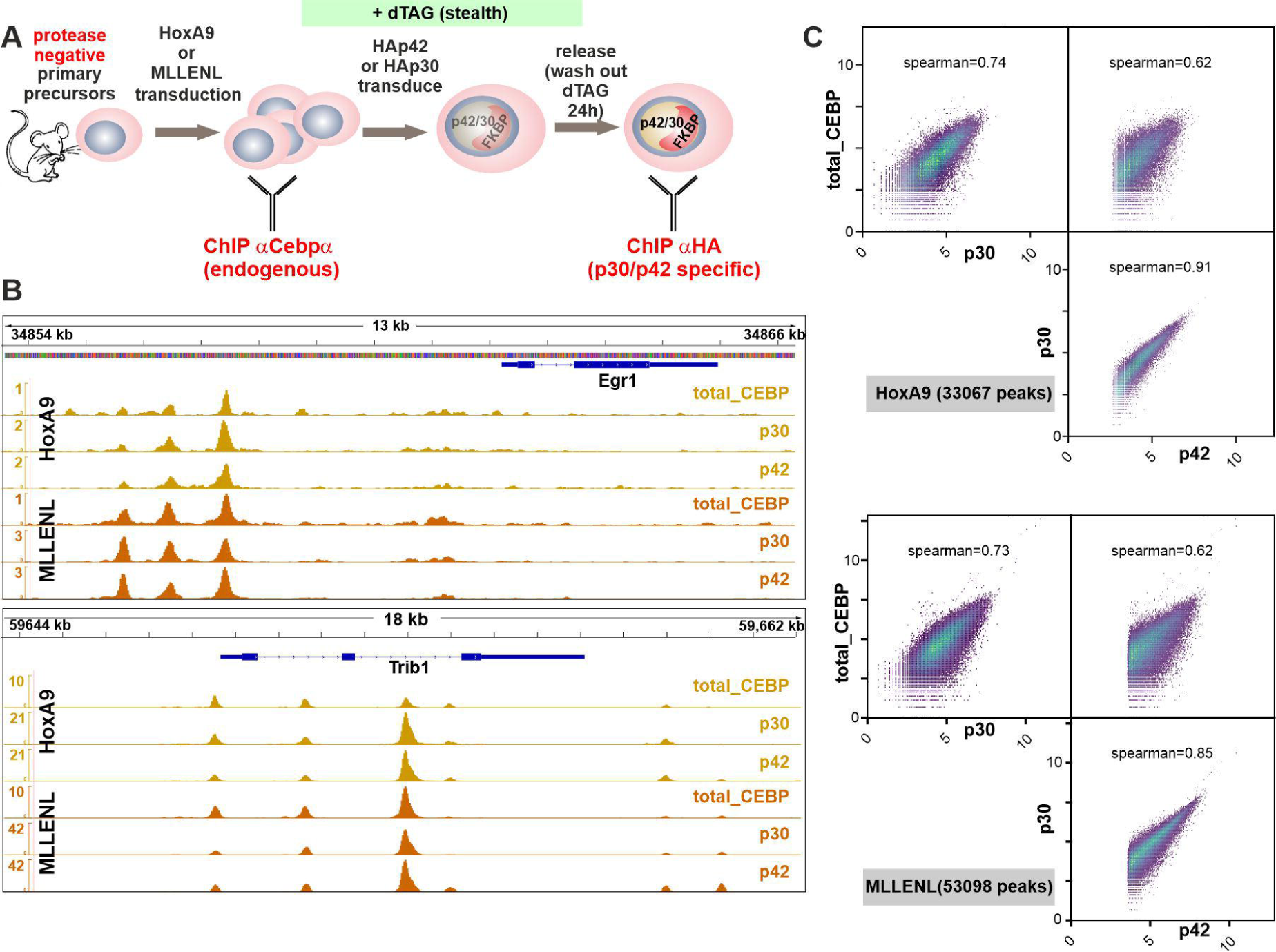
p42 and p30 bind to identical chromosomal loci. A: Schematic overview of experimental setup. B: Integrated genome viewer visualization of p42 and p30 binding in the vicinity of *Egr1* and *Trib1* loci. The graph shows tracks for p42 and p30 as well as for endogenous C/ebpα binding in HoxA9 and MLLENL pre-transformed cells as labeled. C: Global correlation plots of C/ebpα binding. Occupation density of p42/p30/endogenous C/ebpα is plotted against each other at regions identified as p42 peaks in HoxA9 and MLLENL cells as indicated.

### p42 and p30 induce a similar transcriptional program

Next, we wanted to investigate how individual isoforms affect gene expression. As transcription factors primarily affect transcription rates while total RNA amounts are subject to additional controls, we applied nascent RNA sequencing to determine p42 and p30 targets (figure 3A). For this purpose p42 and p30 were induced in HoxA9 and MLLENL cells and nascent RNA was isolated before (0h) as well as 16h and 24h after dTAG-release. Plotting log2-fold changes for both isoforms revealed an overall similar gene regulatory pattern for both isoforms, with the exception of a small group of genes that showed a selective response either to p42 or p30 (supplemental table 1, supplemental figure 3). For further analysis we intersected the gene expression programs of HoxA9 and MLLENL cells and selected common differentially regulated genes with a difference of log2-fold change for p42 versus p30 of 0.75 and greater (for a graphical explanation see figure 3B). Plotting these genes according to their averaged expression across the two cell systems in response to p42/p30 identified a small group of outliers preferentially under control of p42 (figure 3C). Amongst those, two genes *Egr1* and *Trib1* stood out because of their known involvement in leukemia. In humans the *EGR1* (early growth response) gene coding for a transcription factor, is located at 5q right in the center of the genomic region that is deleted in 5q^-^ MDS and AML ^25^. In contrast, *TRIB1* (tribbles1) acts as an oncogene and encodes a protein that has been shown to negatively feedback on C/ebpα ^26^. Screening public databases (www.bioportal.org) confirmed our results, as both genes were significantly underrepresented in AML with *CEBPA* mutations compared to cases with a *CEBPA* wt configuration (figure 3C, inset). Interestingly, in contrast to *TRIB1, EGR1* transcripts were also underrepresented in a considerable number of *CEBPA* wt cases suggesting selective pressure in leukemia to suppress *EGR1* but to retain *TRIB1*.

**Figure 3:**
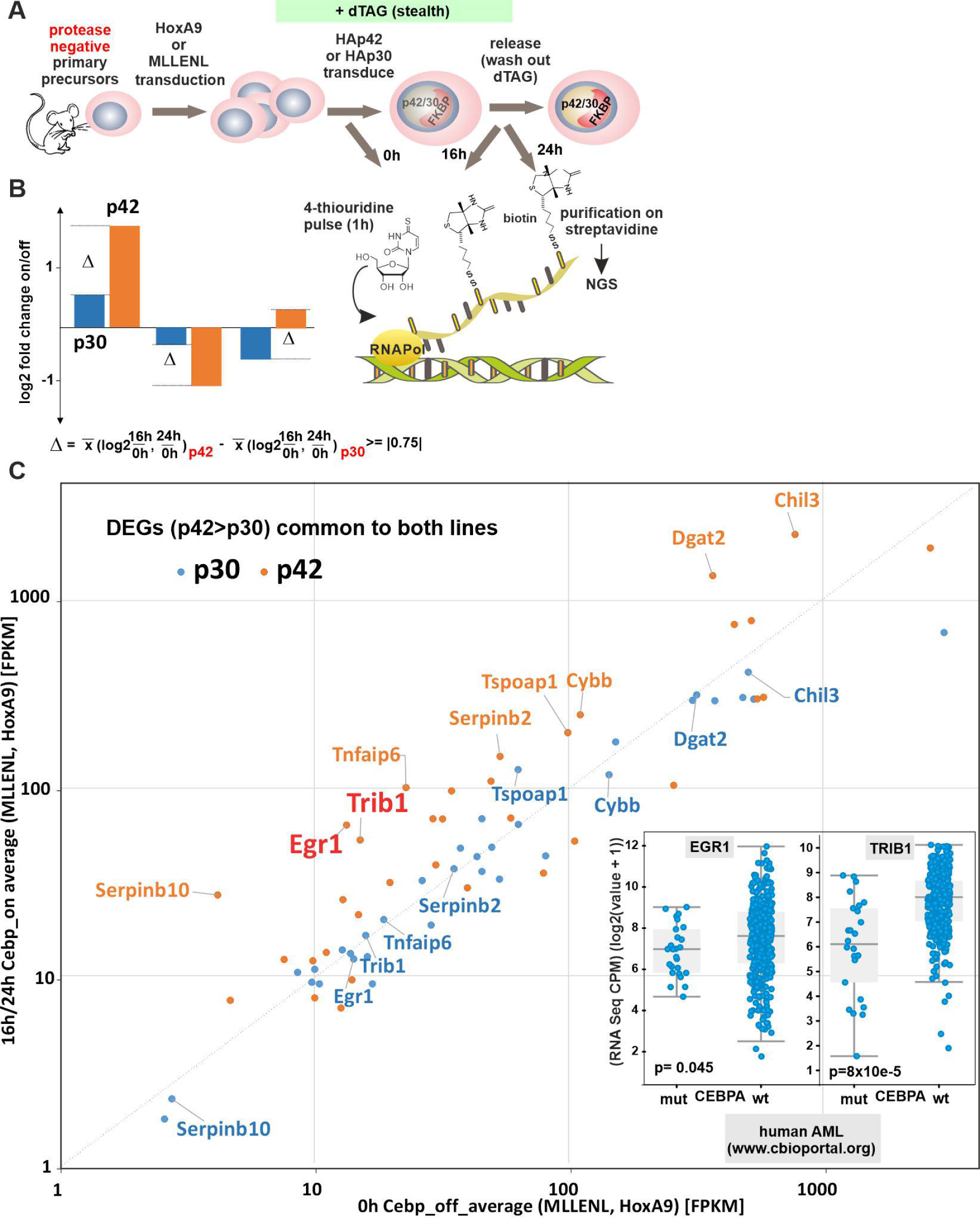
Identification of isotype-specific gene expression. A: Schematic representation of experiment design. At designated time-points cells are labeled for 1h with 4-thiouridine, which is incorporated in newly synthesized RNA. This enables specific purification and NGS-analysis of nascent transcripts. B: Graphical explanation of the criteria applied to designate isotype-specific gene expression. Genes were considered if the Δ between the average log2-fold change between off (dTAG added) and on (dTAG removed at 16h and at 24h) states was larger than 0.75 or smaller than - 0.75. C: Isotype-specific differential gene expression. Only genes fulfilling criteria for differential regulation in HoxA9 and MLLENL cells as above are plotted. Given are average expression levels across both cell lines in the on/off states in FPKM for p42 and p30 expression. Genes with an expression level < 1 FPKM are not considered. Inset: Expression of *EGR1* and *TRIB1* in a human AML cohort as derived from data at www.cbioportal.org. Given are values for *CEBPA* mutant and wt cases.

### Egr1 is a master regulator of myeloid differentiation

To study the physiological implications of Egr1 expression we devised another degron system (figure 4A), because continuous constitutive expression of Egr1 could not be achieved in our myeloid precursors. Indeed, upon release from dTAG mediated degradation Egr1 caused growth arrest similar to p42 (figure 4B). Genomic binding of Egr1 was observed by ChIP mainly at putative enhancer and promoter regions (example in figure 4C). Nascent RNA sequencing of cells after Egr1 induction revealed a gene expression program highly characteristic for myeloid maturation (figure 4D). Most notably the cell cycle inhibitor p21 encoded by *Cdkn1a* was under direct control of Egr1 as well as many other key genes involved in differentiation. Outstanding examples for these were *Id2* (inhibitor of DNA binding) and *Matk.* The Id2 protein displaces E-proteins, which are HSPC-specific bHLH transcription factors from DNA ^27^. Matk is a kinase that induces the nuclear lobulation characteristic for granulocytes ^28^. In GSEA (gene set enrichment analysis), the Egr1 controlled expression program was highly similar to a prototypical myeloid differentiation signature and inverse to a transformation pattern induced by oncogenic HOXA9/MEIS1 (figure 4E).

**Figure 4:**
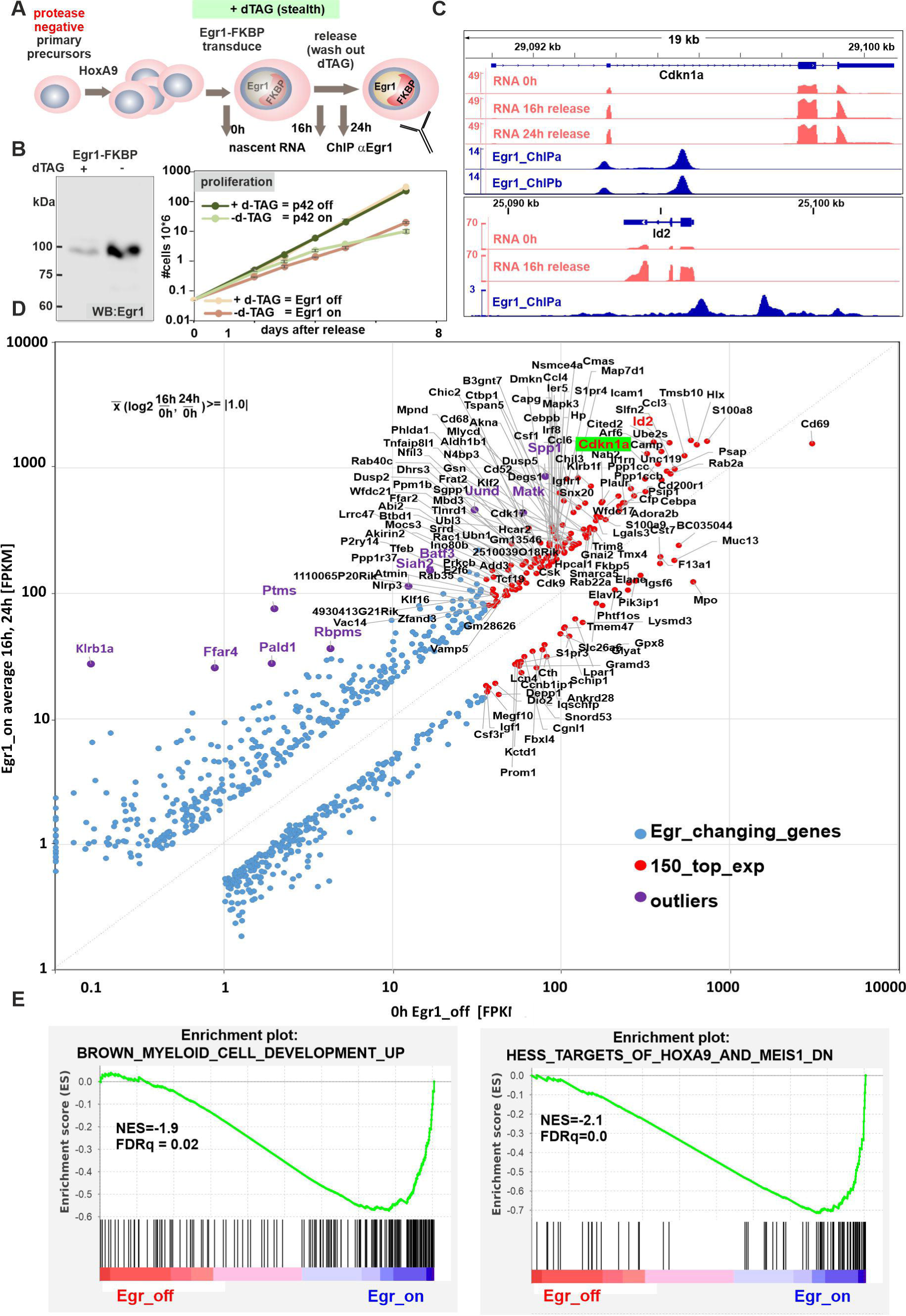
Egr1 is a master regulator of differentiation. A: Schematic overview of genomics experiment. B: Induction of Egr1-FKBP expression in HoxA9 pre-transformed HSPCs induces growth arrest similar to p42. C: Example of a Egr1-specific ChIP and RNA-seq result for select genes *Cdkn1a* and *Id2*. Both ChIP replicates and two timepoints for RNA analysis are given for *Cdkn1a*, only one example each is shown for *Id2*. D: Graphical representation of genes responsive to Egr1. Plotted are average expression values in the Egr1 off and on-states for those genes that showed a log2-fold difference between these conditions of at least 1.0. E: Gene set enrichment analysis of the Egr1-controlled expression pattern.

### Trib1 allows control of differentiation by signaling

Trib1 has been extensively characterized because of its transforming properties. It has been shown that Trib1 binds MEK and induces activation of the MAP-kinase Erk1/2 and it is also involved in a feedback loop curbing C/ebpα activity by inducing specific proteasomal degradation of p42 ^29, 30^. To investigate the paradoxical finding that a transforming gene is amongst the targets of a clearly differentiation-inducing protein, we created Trib1 expressing HSPC lines (figure 5A). In contrast to our experiments with p42 and Egr1, HoxA9 immortalized precursors constitutively expressing Trib1 could be easily produced.

**Figure 5:**
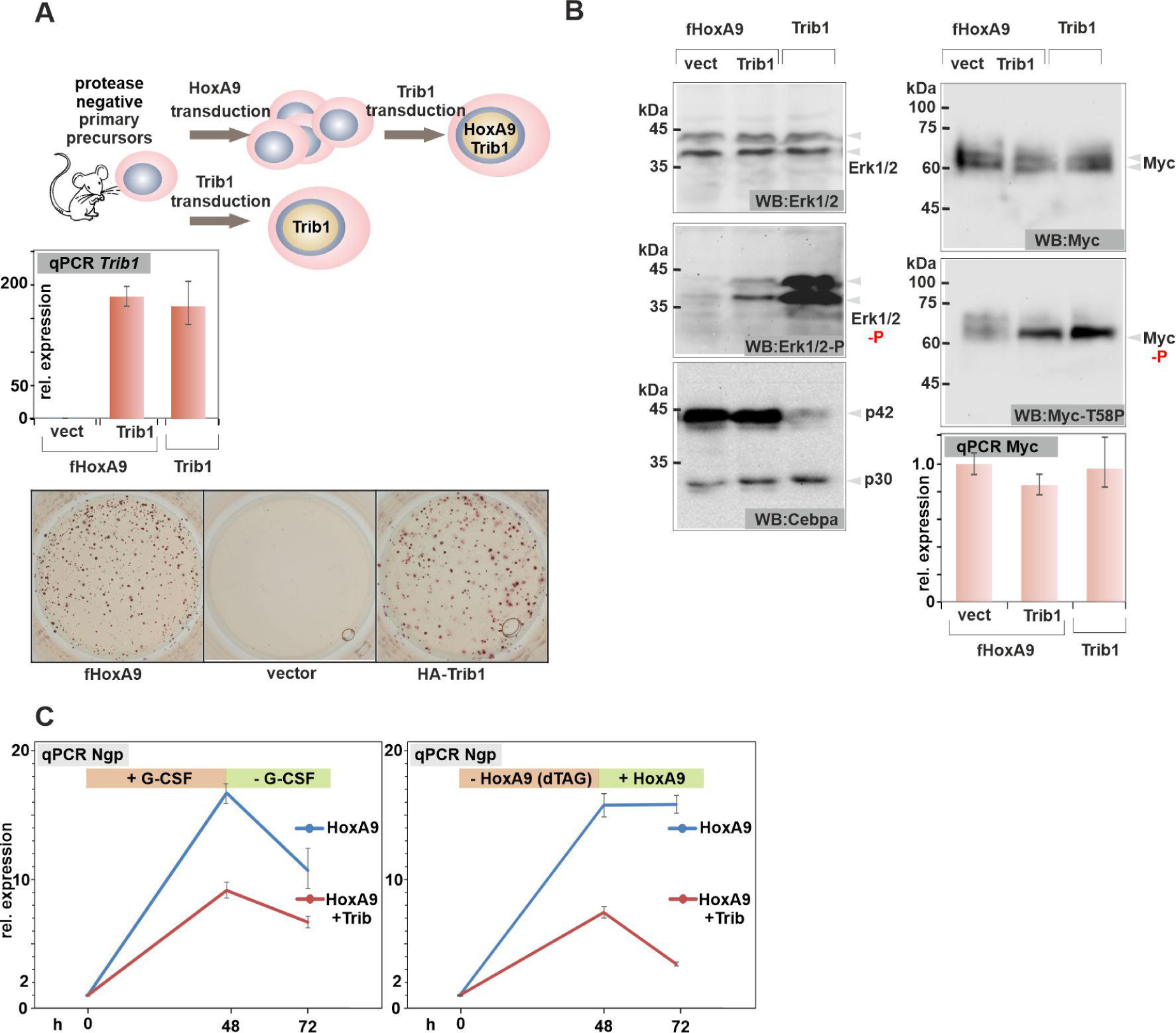
Trib1 blocks differentiation and activates kinase signaling. A: Experimental overview. Trib1 was either added to cells pre-transformed with HoxA9 or it was used for direct transduction of primary HSPCs. As Trib1 protein is highly unstable and cannot be detected by western blot, qPCR was used to confirm successful Trib1 RNA expression as depicted in the bar diagram. The lower panel shows a representative result of a replating assay demonstrating the transforming potential of Trib1 in comparison to HoxA9. B: Trib1 activates Erk1/2 kinases. Western blot experiments with extracts of HoxA9 (control), HoxA9+Trib1, and precursors directly transformed with Trib1 (Trib1) cells. The bar diagram depicts *Myc* RNA expression in the same cells. Immunoblots were developed with antibodies as indicated. C: Trib1 increases resistance against differentiation stimuli. Cells co-expressing Trib1 and HoxA9 (or HoxA9-FKBP) as well as controls were subject to induced differentiation either by cultivation in G-CSF or by degrading HoxA9-FKBP through addition of dTAG for 48h, followed by a 24h recovery period. RNA was isolated at the indicated time-points and expression of the differentiation sentinel gene *Ngp* was determined by qPCR.

Concomitant to published data demonstrating that Trib1 has a short half-live and therefore cannot be detected by conventional immunoblot ^31^, we could not identify Trib1 expression in western blotting, despite considerable over-expression at the RNA level. We also confirmed that it is possible to directly immortalize HSPCs with Trib1 alone. Retroviral transduction of precursors with Trib1 followed by standard replating assays readily yielded immortalized-cell lines. Although cells transformed by Trib1 served well as biochemical controls, they were phenotypically different from the apparently normal myeloid precursor cell lines created by HoxA9 or MLLENL. While Trib1 cells were morphologically myeloid, they were largely devoid of common myeloid surface markers making their classification difficult (supplemental figure 4). Both known activities of Trib1 could be verified. Elevated Trib1 caused increased phosphorylation of Erk1/2 in HoxA9+Trib1 cells and in cells directly transformed by Trib1. A specific reduction of p42 was only observed in the latter cell type (figure 5B). Increased Erk1/2 activity resulted in phosphorylation of Myc a central HoxA9 target protein and a known substrate for Erk-kinases. This modification, however, did not cause a change in Myc protein amount or *Myc* RNA levels. To investigate the impact of Trib1 on differentiation in our cell system, maturation was induced in HoxA9 and HoxA9+Trib1 cells. This was done either by replacing normal cytokines with G-CSF, or by removing the oncogenic stimulus in cells pre-transformed with a degradable HoxA9-FKBP through supplementation with dTAG (see also next paragraph). RNA was isolated before treatment after 48h and 24h after a recovery period where cells were returned to normal growth conditions without G-CSF/dTAG. Expression of the differentiation sentinel gene *Ngp* was determined by qPCR thus allowing a quantitative assessment of differentiation (figure 5C). Trib1 strongly impeded differentiation in both conditions, confirming its pro-transformation activity. Thus, p42 induces Trib1 to introduce a signaling controlled feedback loop to regulate maturation.

### Normal myeloid differentiation intersects downstream of C/ebpα

To further investigate the process of myeloid maturation, we investigated the behavior of HoxA9 immortalized cells during G-CSF/dTAG elicited differentiation. Both regimens induced a maturation program generating morphologically normal granulocytes and monocytes/macrophages (figure 6A). This was accompanied by growth arrest, downregulation of *Myc* and induction of *Ngp*-expression. To examine if this differentiation process is contingent on C/ebpα we followed C/ebpα isoform expression in a time course during treatment (figure 6B). Contrary to expectations, both p30 and p42 expression was rapidly extinguished and there was also no significant shift in isoform ratios. Rather, cell cycle inhibitor proteins p21 and p27 of the Cip/Kip-family increased early after removal of HoxA9 but without detectable Tp53 activation. G-CSF effectively bypassed HoxA9-induced transformation. Concomitant with a previous report ^32^, this was accompanied by a selective upregulation of p27 without affecting p21. These experiments strongly suggest that mechanisms downstream of C/ebpα crucially determine the proliferation/differentiation balance.

**Figure 6:**
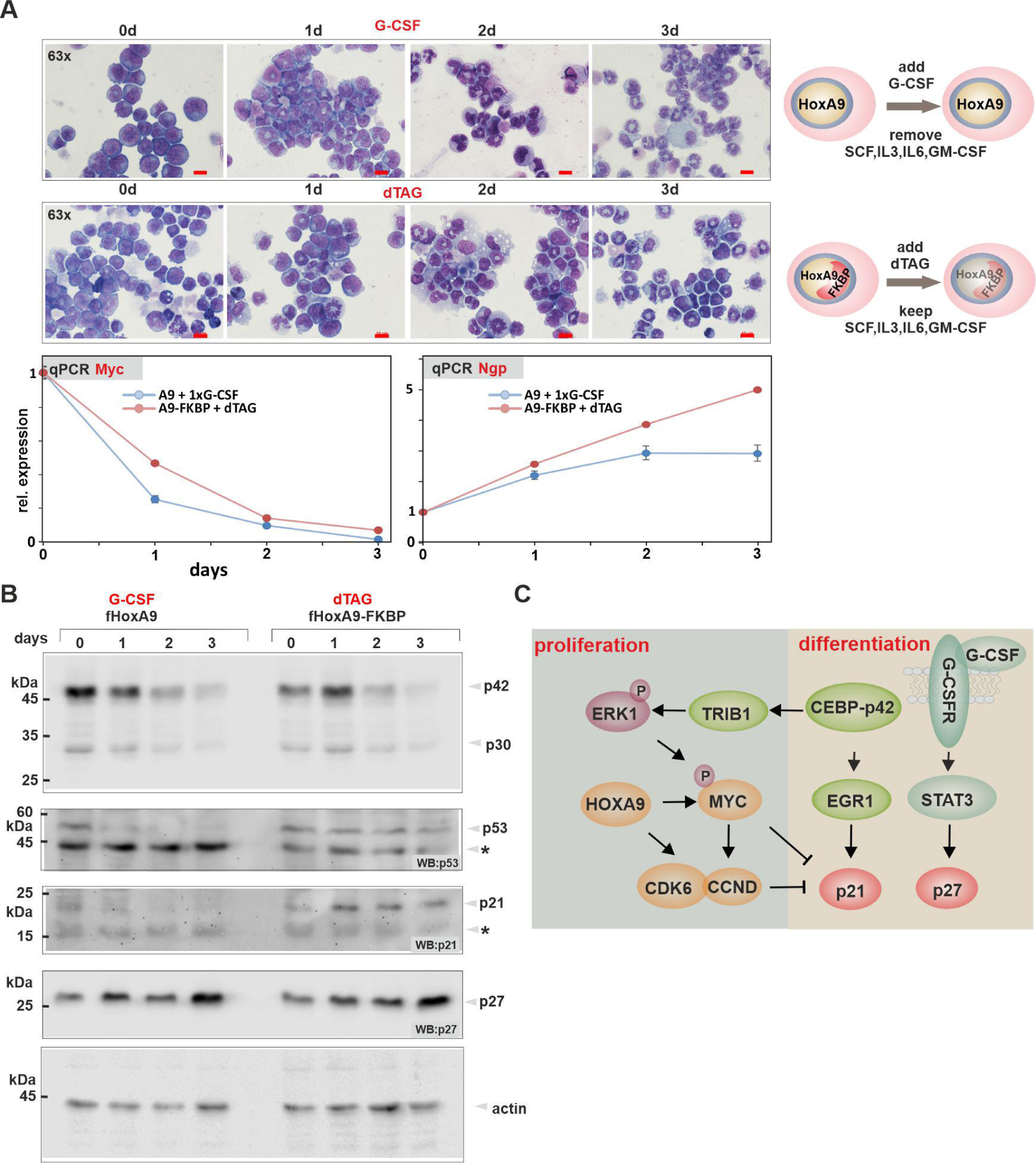
Normal control of differentiation acts downstream of C/ebpα. A: Morphological as well as genomic response of HoxA9/HoxA9-FKBP transformed cells to induced differentiation. Cytospins were stained with May-Grünwald-Giemsa and photographed with a Zeiss Axioskop and a Nikon camera. Shown are 63x magnifications and the scale bar corresponds to 10µM. Gene expression was determined for *Myc* and *Ngp* by qPCR. B: Western blot following endogenous protein expression in a time course during differentiation. Blots were developed with antibodies as indicated. The star denotes unspecific binding. C: Intersection of Hox-mediated transformation with differentiation pathways at the level of the core cell cycle control machinery. The schematic overview shows only part of the known interactions. For further information, please, see text.

## Discussion

Here we show evidence that p42 establishes a “latent” differentiation program that is held in check by signaling. This allows C/ebpα to be expressed in normal precursors without inducing premature maturation. The high pro-proliferative signaling environment during early hematopoietic development enhanced by Trib1 mediated kinase activation enables C/ebpα to cooperate with HoxA9 to establish HSPC specific enhancers without concomitantly pushing the cells into terminal development. Overall, this allows a more precise control of cellular development as would be possible by transcriptional control alone. As our transcriptomics experiments suggest, p30 can functionally replace p42 at most loci. This explains why precursor cells are able to survive with p30 only. The observed mutational pattern of *C/EBPA* in AML creating a functional deletion of p42 while retaining the essential p30 indicates an important role for the loss of function of p42 in cellular transformation. While our results do not exclude a gain of function for the remaining p30, leukemia could well be the consequence of a simple loss of differentiation capacity driven by the absence of p42. This is consistent also with clinical observations as *CEBPA* biallelic mutant AML has a better prognosis in treatment than *CEBPA* wt disease ^33^. In these cases p21 induced by TP53 activation during chemotherapy would substitute for the loss p42/Egr1 activation and bypass the differentiation block similar to experimentally administered G-CSF. Next to cytotoxic effects, neoadjuvant treatment directly elicits differentiation which gives cells with *TP53* mutations an extra survival advantage adding to the dismal prognosis of this type of genetic alteration. Previous reports emphasized the importance of aberrant p30 activity for transformation ^12^. Yet, these studies compared steady state wild-type cells that contained both, p42 and p30 to *Cebpa* mutants with p30 only. Here we manipulate each isoform directly and individually, although with the caveat that this always occurs on top of wt-expression.

Interestingly, HoxA9 did not rely on repression of *Cebpa* to achieve transformation. Rather, proliferation and differentiation pathways seem to intersect further downstream directly at the level of the cell cycle machinery. This is supported by G-CSF induced production of p27 that bypassed transformation directly and induced a state of forced differentiation even in the continuous presence of the transforming event. Generally, the core cell cycle machinery seems to possess a more extended regulatory potential than anticipated. Beyond its role as cell cycle regulator Cdk6 has a nuclear function as transcription factor ^34, 35^ and it is, next to *Myc,* also a central downstream target of HoxA9 ^24^. Together with cyclinD1 whose coding gene *Ccnd1* is strongly activated by Myc, Cdk6/CyclinD1 dimers bind and neutralize p21 protein. In addition, Myc actively suppresses transcription of *Cdkn1a,* the p21 parental gene (figure 6C). Additional cross-connections exist as Cdk6 has been shown to block *Egr1* transcription in conjunction with the transcription factor AP1 ^36^ while active Stat3 in combination with Cdk6 induces *Cdkn2a* ^37^ coding for p16/Ink4a, another cell cycle inhibitor. In summary, these findings point to cell cycle regulators as unexpected major players that coordinate differentiation control by C/ebpα and transforming inputs through Hox-proteins to determine cellular fate.

## Acknowledgements

We thank Renate Zimmermann for technical assistance. This work was supported by research funding from Deutsche Krebshilfe grant 70114166 and in part by the Deutsche Forschungsgemeinschaft grant SL27/9-2 both awarded to RKS.

## Author contributions

MPGC, SA, and RKS performed and analyzed experiments. RKS performed NGS data analysis, conceived and supervised experiments, RKS wrote the manuscript. All authors read and discussed the manuscript.

## Data sharing

NGS reads are available with the European Nucleotide Archive under accession number PRJEB862028.

## Notes

### Competing Interest Statement

The authors have declared no competing interest.

