## Supplemental Data for "A C/ebpα isoform-specific differentiation program in primary myelocytes"

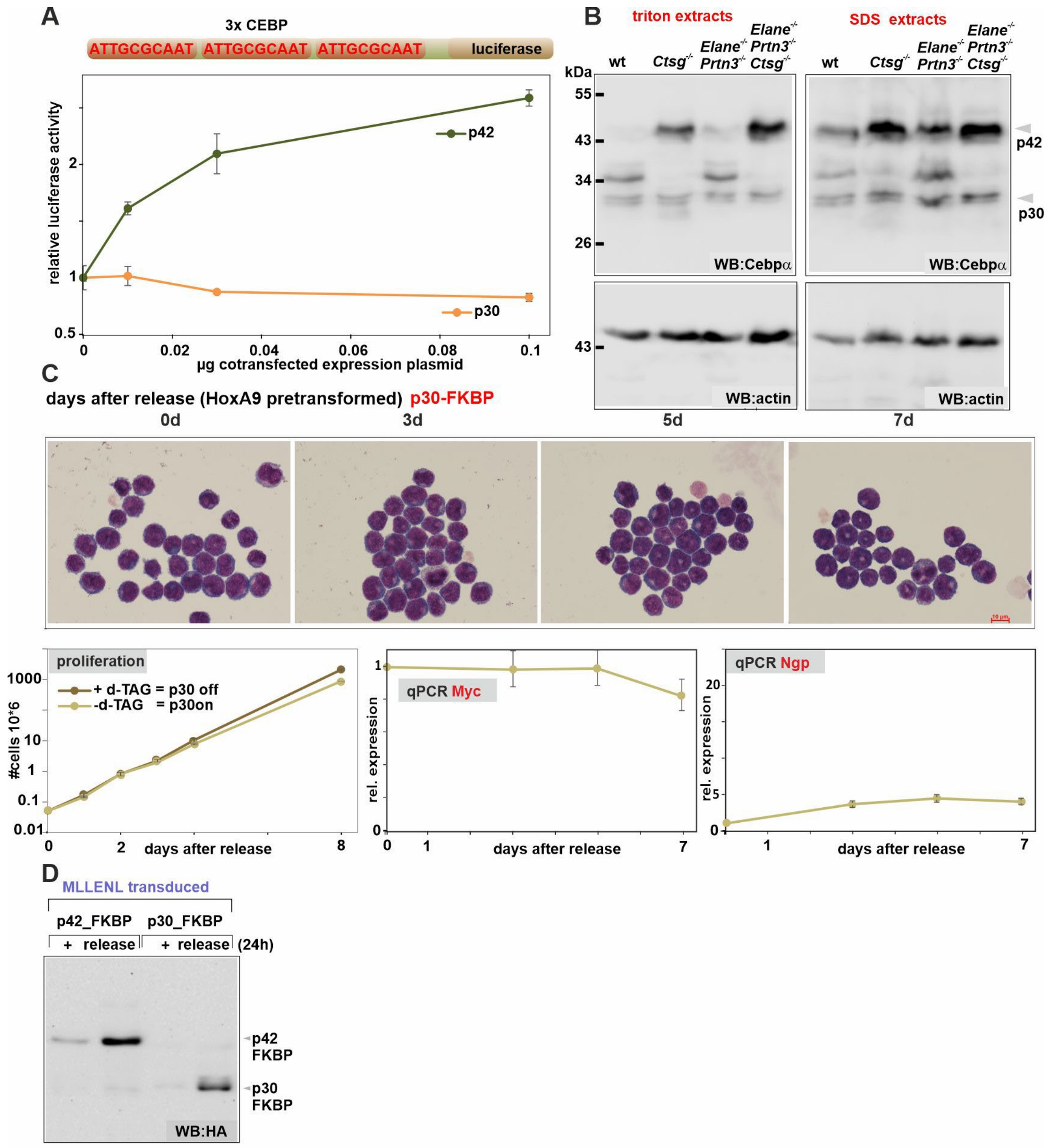


**supplemental figure 1**

### Supplemental Figure 1: Isoform specific properties of C/ebpα

A: p30 is inactive in transient transactivation assays. A luciferase reporter plasmid containing a tandem triplicate of canonical Cebp-binding sites as depicted was used to record transactivation potential of p42 and p30 isoforms. Average and standard deviation of triplicate experiments.

B: p42 is subject to degradation by neutrophil granule proteases. Primary HSPCs from animals of various genetic backgrounds (as indicated) with respect to the presence of granule proteases elastase (Elane), proteinase 3 (Prtn3) and cathepsinG (Ctsg) were transformed with MLLENL. Extracts were generated under lysis conditions comparable to ChIP conditions (triton extracts) and with hot SDS (SDS extracts) to denature all enzymatic activity. Endogenous C/ebpα was detected by western blot.

C: p30 is weakly active in HSPCs. Cells pretransformed by HoxA9 and transduced with p30-FKBP were followed by morphology, proliferation and *Myc*, as well as *Ngp* expression after induction of p30 expression through release from degradation for the indicated time.

D: Expression of p42-FKBP and p30-FKBP in MLLENL pretransformed cells.

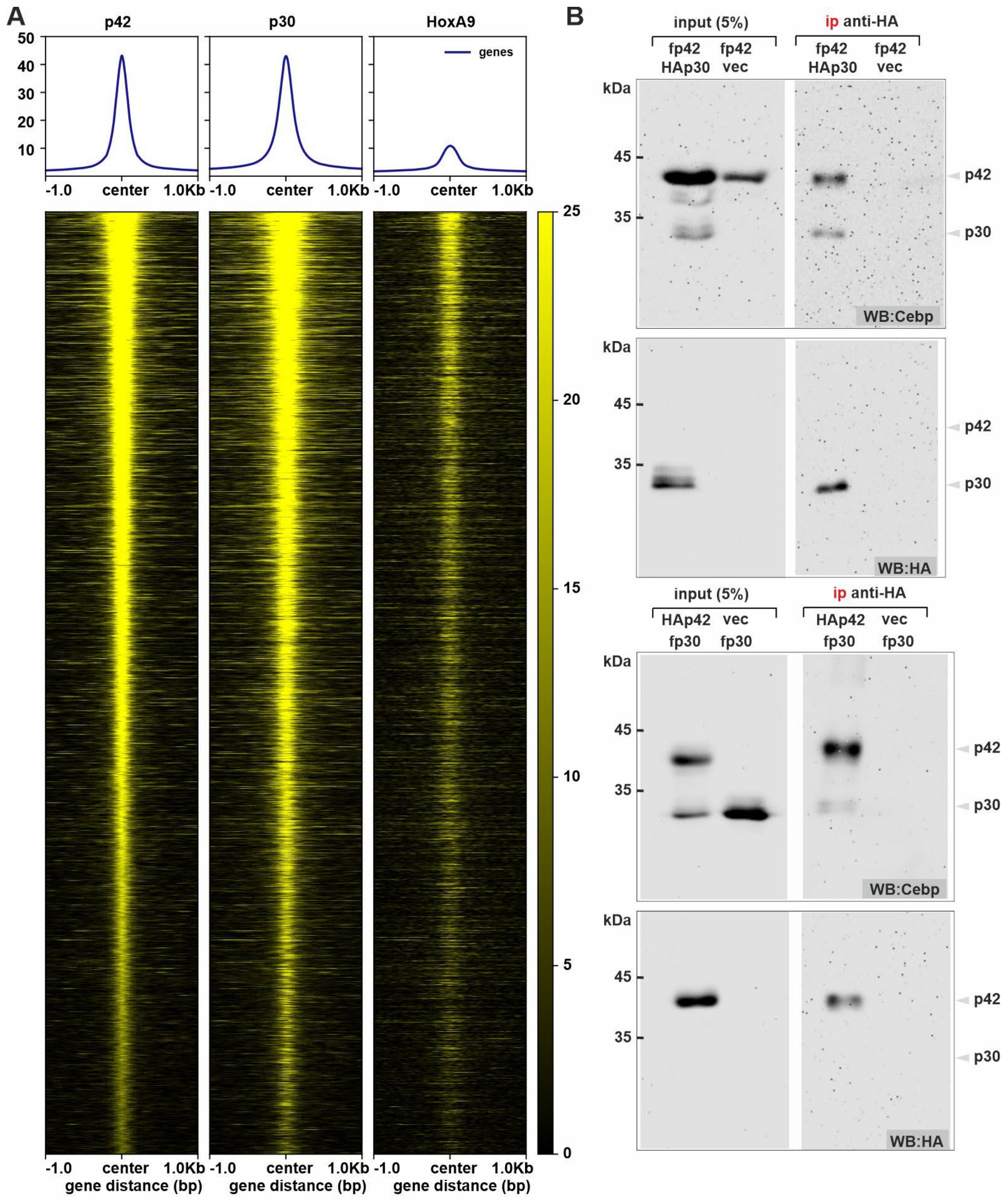


supplemental figure 2

**Supplemental figure 2: Colocalization of p42 and p30 on chromatin**

A: Metaplot of genomic localization of p42, p30 and HoxA9. Binding density of the respective protein is plotted for all 33067 identified p42-peaks in HoxA9 cells. HoxA9 binding data were taken from [1].

B: p42 and p30 directly interact. p42 and p30 N-terminally modified with either flag (f) or HA-tags (HA) were transiently expressed as indicated in 293T cells and immunoprecipitation was done with anti-HA agarose. Westerns were developed with antibodies as labeled.

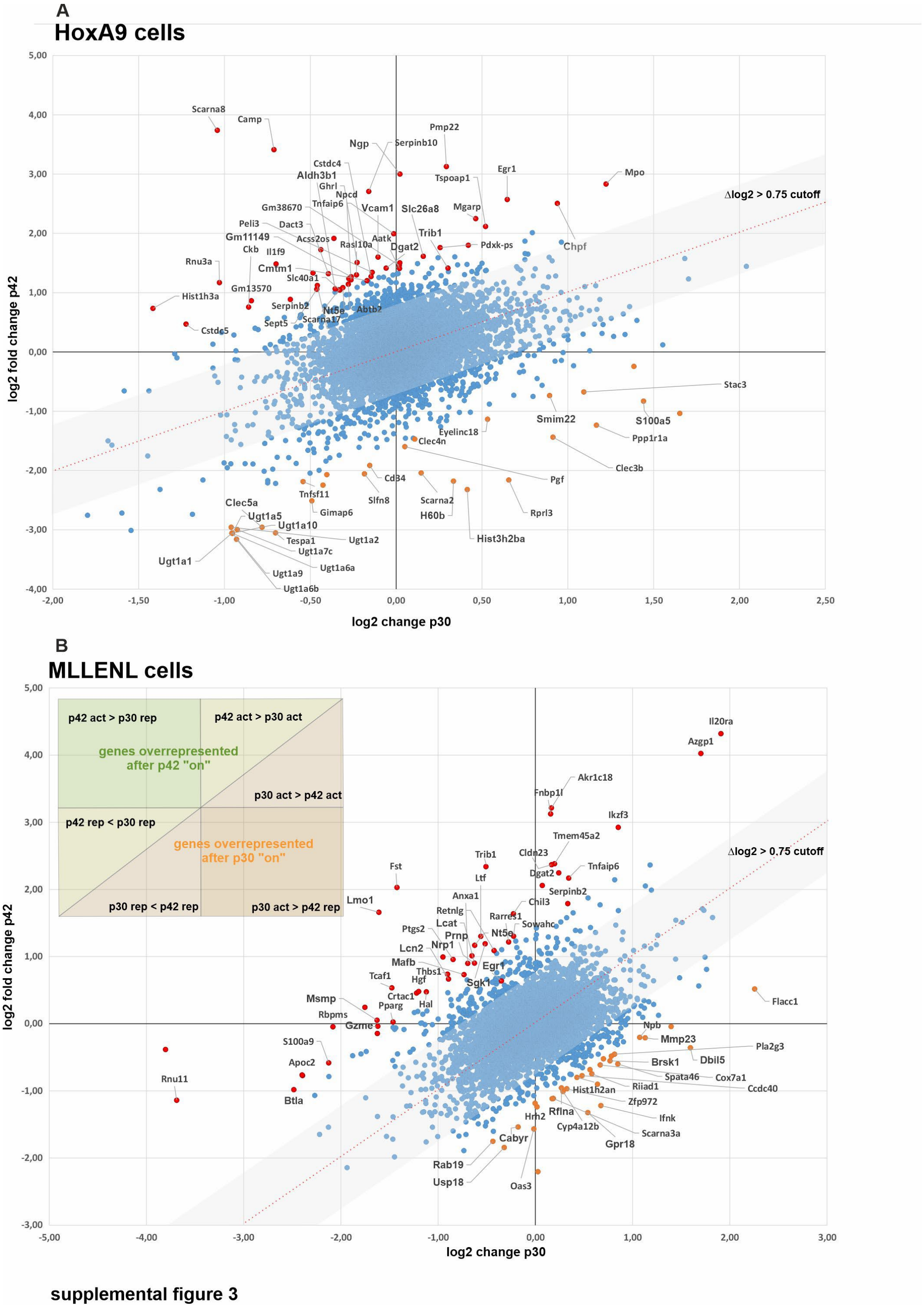


**Supplemental figure 3: The gene expression program governed by p42 and p30.**

A: Effect of p42/p30 expression in HoxA9 pretransformed HSPCs. The average log2-fold change for each gene was calculated from expression ratios of 16h/0h and 24h/0h expression values and plotted for p30 and p42 induction experiments.

B: As for “A” done in MLLENL pretransformed cells.

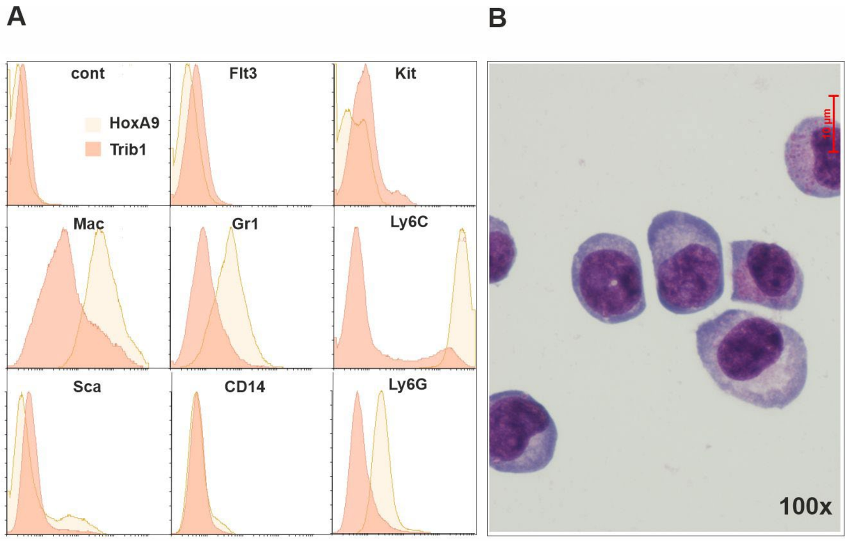


supplemental figure 4

**Supplemental figure 4: Properties of Trib1 transformed primary HSPCs**

A: FACS analysis of surface markers comparing HoxA9 and Trib1 transformed cells as indicated.

B: May-Grünwald-Giemsa stained cytospin of Trib1 transformed cells.

1. Zhong X, Prinz A, Steger J, Garcia-Cuellar MP, Radsak M, Bentaher A, *et al.* HoxA9 transforms murine myeloid cells by a feedback loop driving expression of key oncogenes and cell cycle control genes. *Blood Adv* 2018 Nov 27; **2**(22)**:** 3137-3148.
